# Insecticide resistance and population structure of the invasive malaria vector, *Anopheles stephensi*, from Fiq, Ethiopia

**DOI:** 10.1101/2024.08.21.609030

**Authors:** Jeanne N. Samake, Solomon Yared, Mussie Abdosh Hassen, Sarah Zohdy, Tamar E. Carter

## Abstract

*Anopheles stephensi* invasion in Ethiopia poses a risk of increased malaria disease burden in the region. Thus, understanding the insecticide resistance profile and population structure of the recently detected *An. stephens*i population in Fiq, Ethiopia, is critical to inform vector control to stop the spread of this invasive malaria species in the country. Following entomological surveillance for *An. stephensi* in Fiq, Somali region, Ethiopia, we confirmed the presence of *An. stephensi* morphologically and molecularly in Fiq. Characterization of larval habitats and insecticide susceptibility tests revealed that Fiq *An. stephensi* is most often found in artificial containers and is resistant to most adult insecticides tested (organophosphates, carbamates, pyrethroids) except for pirimiphos-methyl and PBO-pyrethroids. However, the immature larval stage was susceptible to temephos. Further comparative genomic analyses with previous *An*. stephensi populations from Ethiopia using 1,704 biallelic SNPs revealed genetic relatedness between Fiq *An. stephensi* and east-central Ethiopia *An. stephensi* populations, particularly Jigjiga *An. stephensi*. Our findings of the insecticide resistance profile, coupled with the likely source population of Fiq *An. stephensi*, can inform vector control strategies against this malaria vector in Fiq and Jigjiga to limit further spread out of these two locations to other parts of the country and continent.

## Introduction

Malaria is a major global health problem, with an estimated 247 million cases and 619,000 deaths reported in 2021^1^. In Ethiopia, despite past successes in reducing the malaria burden due to the use of indoor residual spraying (IRS) and long-lasting insecticidal nets (LLINs) in malaria control and prevention strategies^2^, it remains a public health concern with an estimated 2 million cases and over 8,000 deaths reported in 2021^1^. The emergence of *An. stephensi*, a malaria vector commonly found in South Asia and parts of the Arabian Peninsula, in the Horn of Africa (HoA)^3-5^, also threatens any gains against malaria in Ethiopia^6^. Indeed, a recent *An. stephensi*-mediated malaria outbreak was reported in Dire Dawa, an urban hub in eastern Ethiopia^7^. Since the detection of *An. stephensi* in the HoA and subsequently, in Sudan (2019)^8^, Nigeria (2020)^9^, Yemen (2021)^10,11^, Kenya (2022)^12^, and Ghana (2023)^13^, the World Health Organization (WHO) has launched an initiative^14^ and released an updated vector alert in 2022^15^ to recommend increased surveillance and research to determine the range of the invasion in order to stop the further spread of *An. stephensi* in Africa. In Ethiopia, *An. stephensi* was first detected in Kebridehar, Somali region, in 2016^4^ and has subsequently been confirmed to be broadly distributed in eastern Ethiopia^16^. However, it was first detected in Fiq, Somali region in June 2021^17^. Malaria cases in the Somali region of Ethiopia are relatively low compared to other regions of Ethiopia ^18^. However, it remains understudied, and more research on malaria parasites and their vectors is needed.

Studies of the breeding sites and the ecology of mosquitoes are very crucial to inform mosquito control strategies such as mosquito larvicidal (temephos) and environmental control (larval habitat removal). Also, the WHO recommends larval source management as one of the immediate control strategies against *An. stephensi* in urban and peri-urban settings of invaded regions^15^. When larval source management or reduction is not feasible, such as a household or town water storage, larviciding may be considered. However, this vector control method can be costly when treating large numbers of larval habitats^19^. Thus, an alternative cost-effective approach is to target specific habitats that can produce large numbers of adult mosquitoes^19^. Thus, determining the Fiq *An. stephensi* susceptibility to larvicide such as temephos can help inform the decision making for this control method against the invasive malaria vector in Fiq town.

Moreover, genomic analyses can further inform control strategies against the newly detected Fiq *An. stephensi*. Specifically, assessing the genetic diversity and population structure of Fiq *An. stephensi* population in comparison to an established one in the region can give insights into its demographic history, dispersal patterns, and potential source populations.

Hence, we conducted an entomological survey after a year of the first detection of *An. stephensi* in Fiq town, Somali region, Ethiopia to first characterize larval habitats for *An. stephensi* and determine their insecticide susceptibility status, including susceptibility to the larvicide, temephos. After identifying the morphology, we confirmed it molecularly and used genomic approaches to analyze the demographic history and population structure of *An. stephensi* in Fiq. We compared the population structure with the previously detected *An. stephensi* populations in eastern Ethiopia to determine the extent of establishment in Fiq. We further assessed their genetic connectivity to those populations to uncover their potential source populations within the region.

## Material and Methods

### Study setting

Fiq is a small town located in the Somali region of Ethiopia. It is situated at an altitude of 1200 m and 195 km west of Jigjiga, the capital city of the Somali Region. The town features sporadic mountains and small house constructions. It is common to see water stored in pits for building bricks for house construction.

### Sample collection

Entomological survey was carried out in Fiq town during the rainy season from May to June 2022, by Jigjiga University. Adult mosquitoes were collected indoors and outdoors using pyrethrum spray collections (PSC) and CDC light traps (CDC LT), respectively, and Prokopack aspiration from animal shelters. Sampling of mosquito larvae was conducted from houses with water storage containers, particularly, cisterns and plastic sheet water storage. During each survey, a habitat was first visually inspected for the presence of mosquito larvae, and then twenty samples were taken with a soup ladle (350 ml capacity) from each breeding habitat (see Supplementary Fig. S1 online). The *Anopheles* larvae were separated from the Culicine larvae based on their position to the water surface. Collected *Anopheles* larvae were reared to the adult at a field laboratory for morphological species identification (see Supplementary Fig. S2 online).

### Mosquito identification

Adult *Anopheles* were morphologically identified at the species level using the Afrotropical mosquitoes key^20^ (see Supplementary Fig. S2 online). Polymerase chain reaction (PCR) assays were conducted on a subset of *An. stephensi* specimens identified by morphology. These assays targeted portions of the mitochondrial cytochrome oxidase subunit I (*COI*) and the nuclear internal transcribed spacer 2 (*ITS2*) loci for molecular identification, as previously reported^4^. For further confirmation of species identification, we performed phylogenetic analysis with the generated *COI* sequences and those of previously detected *An. stephensi* from Ethiopia retrieved from GenBank (Accession# OK663483, OK663480, OK663484, OK663481, OK663479, OK663482)^21^. These accession numbers represent unique *COI* haplotypes of *An. stephensi* previously identified across eastern Ethiopia^21^. Phylogenetic relationships were inferred using a maximum likelihood approach with RAxML GUI^22^ using the GTR model of nucleotide substitutions, gamma model for rate of heterogeneity (GTRGAMMA option), and one thousand replicates in one run for bootstrap analysis. *Anopheles maculatus* was designated as an outgroup. The tree with the highest log likelihood was visualized and formatted in FigTree^23^.

### WHO insecticide susceptibility tube tests

Insecticide susceptibility tests were conducted on adult female *An. stephensi* reared from wild larvae (see Supplementary Fig. S2 online) following standard procedures^24^. One hundred mosquitoes from each population were tested for each insecticide using the diagnostic concentration, and 50 mosquitoes were used for controls. The insecticides used were 0.1% bendiocarb, 0.25% pirimiphos-methyl, 0.05% alpha-cypermethrin, 0.05% deltamethrin, 0.75% permethrin, and 0.15% cyfluthrin. Based on the WHO mortality criteria, resistance was determined as follows: 98–100% mortality indicates susceptibility, 90– 97% mortality indicates possible resistance (requires further investigation), and less than 90% mortality confirms resistance ^19^.

### Piperonyl butoxide synergist assays

Piperonyl butoxide (PBO) synergist assays were conducted on *An. stephensi* against two pyrethroids (deltamethrin and permethrin). The synergist assays were conducted by pre-exposing mosquitoes to a 4% PBO paper for 60 min. Mosquitoes were then transferred to tubes with the pyrethroid of interest for 60 min and the susceptibility was determined based on the WHO mortality criteria^24^ described above.

### Temephos susceptibility tests

Larval susceptibility tests were performed to assess the sensitivity of *An. stephensi* larvae to temephos, an organophosphate larvicide. According to WHO procedure^25^, conventional techniques were employed for the testing, and 312.5 mg/l, 62.5 mg/l, 12.5 mg/l, and 2.5 mg/l concentrations were utilized to generate a final concentration of 1.25mg/l, 0.25mg/l, 0.05mg/l, and 0.01mg/l when 1 ml of each concentration was added to 249 ml of normal water. To detect resistance, an estimated diagnostic dosage of 0.25 mg/l was employed^26^, and larvae mortality data were interpreted following the same WHO mortality criteria used for adult mosquitoes^19^. For each larvicide concentration, four duplicates of 25 larvae were employed, with two replicates serving as controls. To get 100 larvae examined per dosage, four beakers were utilized per dose. After 24hrs all larvae without movement on the water surface were considered as dead.

### Target-site insecticide resistance loci analysis

To analyze phenotypic and genotypic associations in observed pyrethroid-resistant *An. stephensi*, we sequenced the pyrethroid target site in the voltage-gated sodium channel (*vgsc*) gene and downstream intron to genotype any knockdown resistance mutation (*kdr*). Analyses were conducted using previously published protocols^27,28^. To genotype any *kdr* mutations, sequences were queried to NCBI BLAST to confirm correct *kdr* locus, then aligned to reference sequences from Samake et al.^27^ using CodonCode Aligner version 8 (CodonCode Corp., Centerville, MA, USA). We then generated an updated Ethiopian *An. stephensi kdr* mutation phylogenetic tree with *kdr* mutation sequences from Samake et al.^27^ also including the only non-African *An. stephensi kdr* mutation sequence available in Genbank (Accession# JF304952)^28^. Phylogenetic relationships were inferred using a maximum likelihood approach with RAxML GUI^18^ using the GTR model of nucleotide substitutions, gamma model for rate of heterogeneity (GTRGAMMA option), and one thousand replicates in one run for bootstrap analysis. The tree with the highest log likelihood was visualized and formatted in FigTree^23^.

### Nuclear population structure and genetic diversity

To assess nuclear population structure and potential source populations, we first generated double digest restriction-site associated DNA (ddRAD) sequences of Fiq *An. stephensi* in addition to previously published raw ddRAD sequences of Ethiopian *An. stephensi* ^29^. Genomic DNA was extracted from adult *An. stephensi* mosquitoes using Qiagen DNeasy Blood and Tissue kit (Qiagen). DNA quality was assessed on a 1% agarose gel to ensure at least 10 kilobase DNA fragments and quantified using Nanodrop One Spectrophotometer (Thermo Fisher Scientific Inc.) to ensure a minimum concentration of 20 ng/μl. ddRAD-seq library preparation followed protocols outlined in Lavretsky et al.^30^ (also see Samake et al.^29^). In short, each genomic DNA was enzymatically fragmented using SbfI and EcoRI restriction enzymes and ligated with Illumina TruSeq compatible barcodes for demultiplexing purposes. Libraries were quantified using Qubit dsDNA BR Assay Kit (ThermoFisher Scientific, MA, USA), pooled in equimolar amounts, and sequenced using single-end chemistry sequencing on an Illumina HiSeq X at Novogene (Novogene CO., Ltd., Sacramento, CA, USA; see detailed methods in Supplementary Document S1 online). We then retrieved raw sequence reads of previously reported *An. stephensi* populations from 10 different sites across eastern Ethiopia (n = 183, BioProject PRJNA888109, Samake et al.^29^) to generate a combined bi-allelic SNPs dataset. Raw Illumina reads were demultiplexed, processed and SNP genotyped using the computational pipeline described in Lavretsky et al.^31^ (also see Samake et al.^29^). We used Trimmomatic ^32^ to trim or discard poor-quality sequences using a Phred score of ≥30 to ensure only high-quality sequences were retained. We then used the Burrows-Wheeler Aligner^33^ (bwa) to align the remaining quality reads to the *An. stephensi* reference genome (Accession PRJNA661063; Chakraborty et al.^34^). Samples were sorted and indexed in Samtools and genotyped using the ‘mpileup’ function in BCFtools ^35^.

We then identified variation among samples with a principal component analysis (PCA) using the --pca function in PLINK v.1.9 ^36^ and visualized with the R package ggplot2^37^. Next, individual maximum-likelihood population assignment probabilities were attained across samples using ADMIXTURE v.1.3^38^. Each ADMIXTURE analysis was run with a 10-fold cross validation (CV) and with a quasi-Newton algorithm to accelerate convergence^39^. To limit possible stochastic effects, each analysis was based on 1,000 bootstraps for each population *K* value of 1-10. The block relaxation algorithm for point estimation was used for each analysis and terminated once the log-likelihood of the point estimation increased by <0.0001. The optimum population value was based on the average of CV-errors across the analyses per *K* value. ADMIXTURE assignment probability outputs were visualized using the R package ggplot2^37^. Additionally, nucleotide diversity (π), counts of segregating SNPs (S), Tajima’s D, and pairwise estimates of fixation index (F_st_) by site were calculated in the R package PopGenome^40^.

### Genetic network

To further access potential source populations of the Fiq *An. stephensi* population, we performed a network analysis with the combined biallelic SNPs dataset from Fiq sequences (n = 20) and Genbank retrieved *An. stephensi* sequences from 10 different sites across eastern Ethiopia (n = 183, Samake et al.^29^). We used EDENetworks^41^, which allows network analyses based on genetic distance matrices without a prior assumption. The network consists of nodes representing populations connected by edges/links weighted by their F_st_ based Reynolds’ genetic distances (D)^42^ which provide the strength of connectivity between pairs of populations^41^. The thicker the edge/link, the stronger the genetic connectivity between the two populations. Moreover, node size is proportional to the cumulative weighted edge linkages for each population. Thus, the larger the node the higher the connectivity hub or sink. Statistical confidence of the nodes was evaluated using 1,000 bootstrap replicates. Nodes that appear in the top 5 and top 1 lists of betweenness centrality (BC) values (number of shortest genetic paths passing through a node) can be considered as statistically significant^43^.

## Results

A total of 221 adult mosquitoes were collected using CDC LT, PSC, and Prokopack from animal shelters. Of these, 219 adults *Culex* spp, and only two adult *An. stephensi* were collected using Prokopack aspiration (see Supplementary Table S1 online). No other *Anopheles* mosquitoes were collected using PSC and CDC LT.

### *Anopheles stephensi* larval habitats

Thirty-one mosquito potential breeding sites were inspected. Twenty-six containers including eight plastic ones, fifteen cisterns, and three barrels were positive for mosquito larvae (see Supplementary Table S2 online). There were no mosquito larvae recovered from five containers. A total of 4103 larvae of mosquito were collected from these different breeding sites. Of these, 3713 *Anopheles* and 390 *Culex* were identified (see Supplementary Table S2 online). All of the *Anopheles* mosquitoes were reared to adults successfully and morphologically identified as *An. stephensi*.

### Insecticide susceptibility and synergist assays

According to the WHO bioassay procedure, 1200 non-fed female *An. stephensi* were tested with various insecticide classes and PBO-pyrethroids. Based on the WHO mortality criteria, Fiq *An. stephensi* revealed resistance to all insecticides tested, except for pirimiphos-methyl which resulted in 100% mortality (Figure 1a; also see Supplementary Table S3 online). Pre-exposure to PBO restored complete sensitivity to deltamethrin and permethrin in the synergist tests (Figure 1a; also see Supplementary Table S3 online).

**Figure 1.**
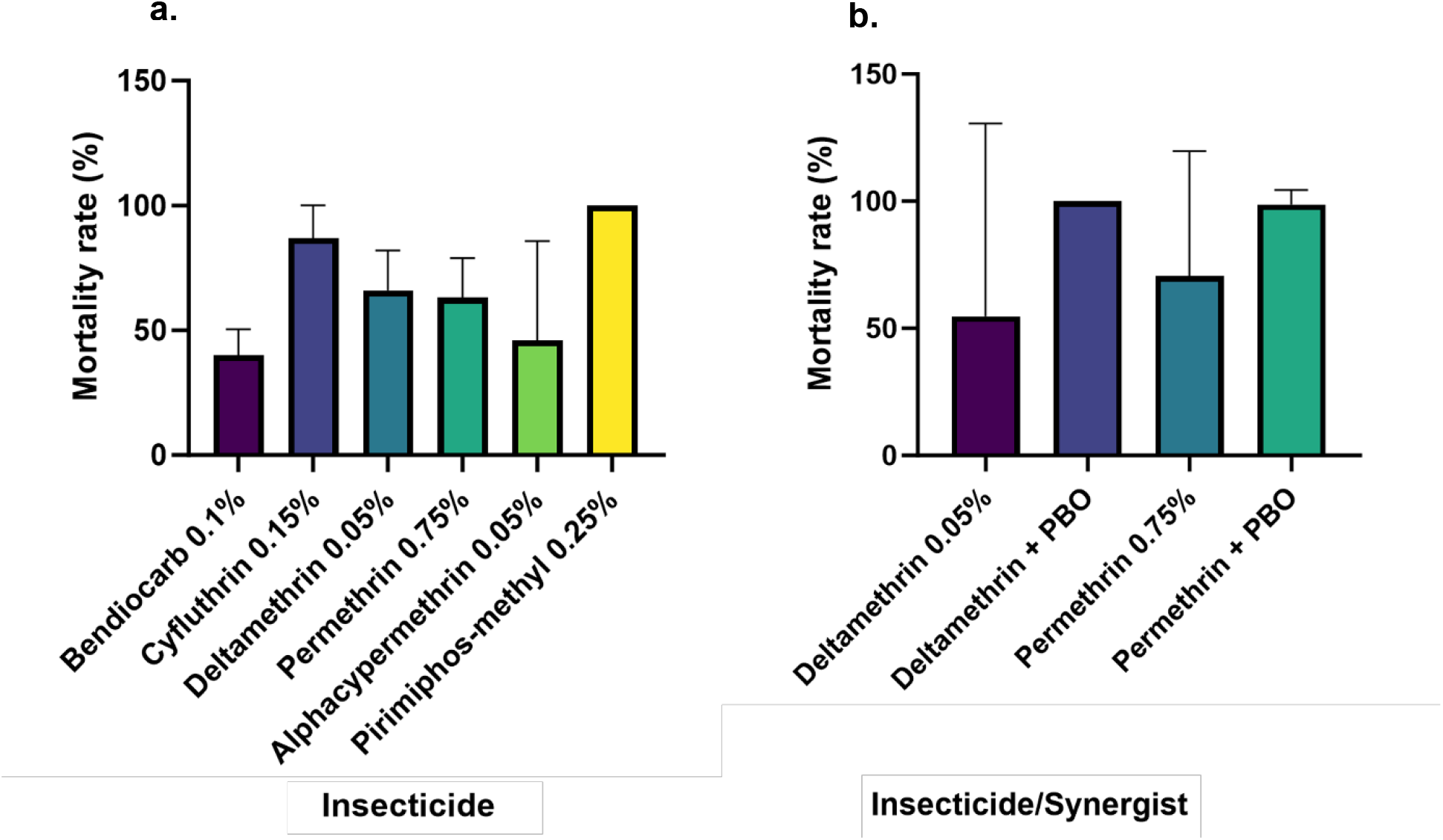
Mortality rate of *An. stephensi* exposed to different insecticides in Fiq, Ethiopia. Bars represent mean values (with 95% CIs) **a**. Diagnostic dose assays. **b**. Synergist assays.

### Temephos susceptibility

The Temephos susceptibility test against Fiq *An. stephensi* larvae revealed that concentrations of 1.25mg/l and 0.25mg/l killed 100% of the *An. stephensi* larvae after 24hrs (Table 1). However, mortality rates for the rest of the concentrations such as 0.05mg/l and 0.01mg/l were 94% and 16%, respectively, below the WHO mortality criteria of greater than 98 percent for susceptibility (Table 1).

**Table 1.** Mortality rate of *An. stephensi* larvae against WHO standard concentrations of Temephos.

| Concentration | Mortality rate (%) | Mortality rate (%) |
| --- | --- | --- |

|  | Final<br>concentration | R1 | R2 | R3 | R4 | mean [95% CI] |
| --- | --- | --- | --- | --- | --- | --- |
| 312.5mg/l | 1.25mg/l | 100 | 100 | 100 | 100 | 100 |
| 62.5mg/l | 0.25mg/l | 100 | 100 | 100 | 100 | 100 |
| 12.5mg/l | 0.05mg/l | 100 | 96 | 80 | 100 | 94 [79, 109] |
| 2.5mg/l | 0.01mg/l | 12 | 8 | 20 | 24 | 16 [4, 28] |

NB: A total of 1000 adult *An. stephensi* identified and no other *Anopheles* species encountered to provide proof that the susceptibility test was done on *An. stephensi* larvae. R1-R4 are replicates.

### Molecular identification

A subset of 154 morphologically identified *An. stephensi* collected from the various sites in Fiq were preserved with silica gel and sent to Baylor University for molecular and genomics analyses. Genomic DNA was extracted from 20 of those specimens for a comparative sample size with previous *An. stephensi* SNPs dataset from eastern Ethiopia^29^. The 20 samples were successfully confirmed as *An. stephensi* based on the presence of the characteristic band in the ITS2 endpoint assay (see Supplementary Fig. S3 online), and phylogenetic analysis of the cytochrome oxidase subunit I (COI) mitochondrial marker further confirmed *An. stephensi*. All the analyzed Fiq *An. stephensi* COI sequences clustered with the most prevalent Ethiopian *An. stephensi* COI haplotype (Hap2)^21^ (see Supplementary Fig. S4 online).

### *Kdr* mutation population frequency

Of the 20 *An. stephensi* samples analyzed for pyrethroid target site phenotypic-genotypic association in observed resistant *An. stephensi*, one (5%) carried the *kdr* L1014F mutation with a heterozygote allele (see Supplementary Table S5 online). The *Kdr* L1014S was not observed. Phylogenetic analysis confirmed the Fiq *kdr* L1014F mutation, as the analyzed sequence clustered with previously published *An. stephensi kdr* L1014F mutation sequences from Ethiopia^27^ and India^28^ (bootstrap 100) (Figure 2).

**Figure 2.**
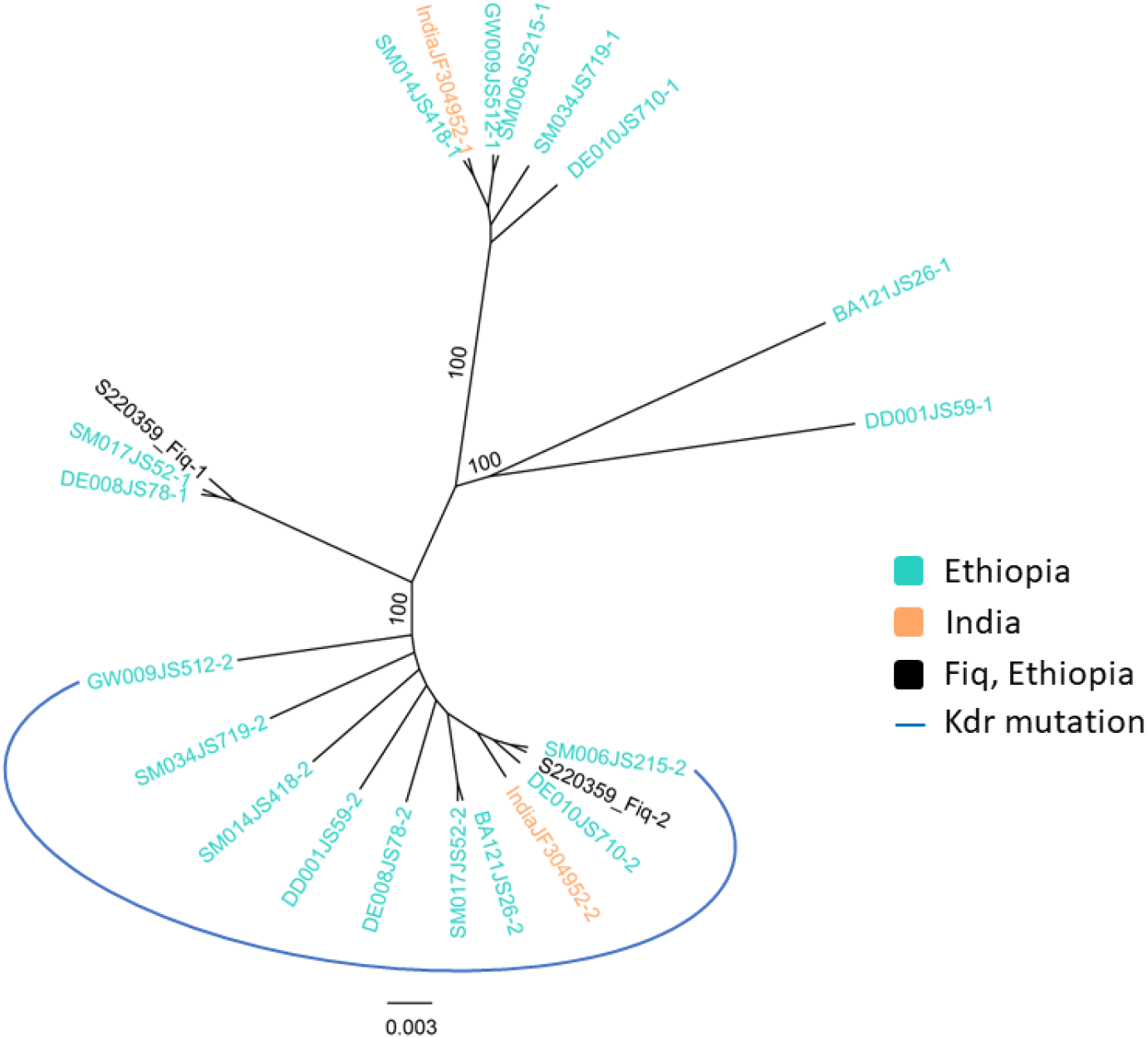
Phylogenetic tree of *kdr* L1014F mutation in Fiq *An. stephensi* and available global *An. stephensi kdr* mutation sequence. The evolutionary history was inferred by using the Maximum Likelihood method based on the General Time Reversible model. The tree with final ML optimization likelihood (−146.34) and bootstrap values >70 are shown.

### Nuclear genetic diversity and population structure

For a comparative analysis between the Fiq studied population and the previously reported *An. stephensi* populations in 10 different sites across eastern Ethiopia^29^, genetic diversity and population structure assessments were based on a combined SNPs dataset of 1,704 independent bi-allelic SNPs after filtering for linkage disequilibrium and non-biallelic SNPs. From the combined Ethiopian *An. stephensi* bi-allelic SNPs analyzed, the lowest nucleotide diversity was observed in Godey (π = 0.1979) in southeastern Ethiopia, and the highest nucleotide diversity was observed in Awash Sebat Kilo (π = 0.2300) in northeastern Ethiopia (see Supplementary Table S4 online). Fiq had the second lowest nucleotide diversity (π = 0.1986) (see Supplementary Table S4 online). *Anopheles stephensi* populations from southeastern Ethiopia and Fiq were also found to have the highest Tajima’s D values, 1.36, 1.48, 1.19, and 1.29 for Degehabur, Kebridehar, Godey, and Fiq, respectively, indicating a lack of rare variants relative to neutral expectations^44^ (see Supplementary Table S4 online).

Assessing population structure using the combined *An. stephensi* independent bi-allelic SNPs dataset and based on principal component analysis (PCA), we identified three semi-discrete genetic clusters (Figures 3b and 3c). Plotting the first two principal components grouped *An. stephensi* populations regionally with Fiq *An. stephensi* mostly clustering with east central Ethiopia *An. stephensi* populations (Figure 3). The ADMIXTURE analysis based on an optimum population K model 5 more specifically identified five main genetic similarities across *An. stephensi* populations that include (1) Erer Gota, Dire Dawa, and Jigjiga, (2) Fiq, (3) Bati and Semera, (4) Gewane and Awash Sebat Kilo, and (5) Degehabur, Kebridehar, and Godey (see Supplementary Fig. S5 online). From these genetic clusters, Fiq *An. stephensi* displayed the lowest admixture proportions as it is composed of a single ancestral lineage that is predominantly observed in Jigjiga admixed *An. stephensi* population (see Supplementary Fig. S5 online). The F_st_ pairwise estimates also showed Fiq *An. stephensi* being less differentiated from Jigjiga *An. stephensi* from east central Ethiopia populations (F_st_ = 0.07) and more differentiated from Bati *An. stephensi* from northeastern Ethiopia populations (F_st_ = 0.14) (see Supplementary Fig. S6 online).

**Figure 3.**
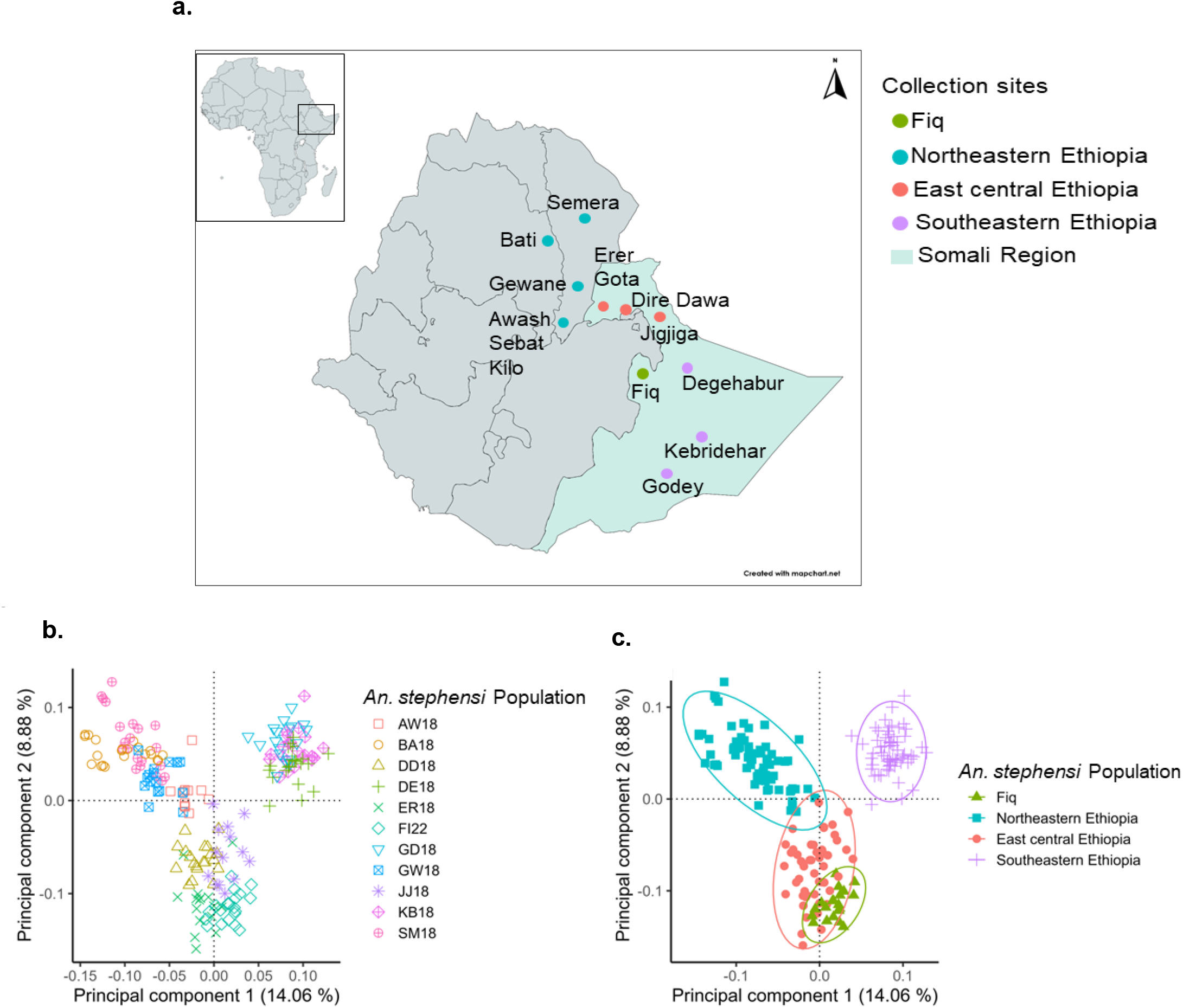
**a**. *Anopheles stephensi* sampling locations in eastern Ethiopia. Fiq data was generated in this present study. Data from all other sites were previously generated (Samake et al.^29^, 2023). Map was created with mapchart.net. **b**. Principal component analysis (PCA) of individual *An. stephensi* population in eastern Ethiopia. The amounts of variation explained by each principal component (PC 1 on the x-axis, PC 2 on the y-axis) are given in percentages. The sample sites are Awash Sebat Kilo (AW18), Bati (BA18), Dire Dawa (DD18), Degehabur (DE18), Erer Gota (ER18), Fiq (FI22), Godey, (GD18), Gewane (GW18), Jigjiga (JJ18), Kebriderar (KB18), Semera (SM18). **c**. Scatterplot of subgroup variation based on principal component analysis (PCA). The amounts of variation explained by each principal component (PC 1 on the x-axis, PC 2 on the y-axis) are given in percentages. *Anopheles stephensi* population subgroups are color-coded as described in the legend.

### Genetic network

Network reconstruction was based on the same combined 1,704 independent bi-allelic SNP dataset. Out of the ten Ethiopian *An. stephensi* populations analyzed, Fiq *An. stephensi* was found to be connected to two populations, Dire Dawa and Jigjiga, that were recovered as statistically significant nodes based on the bootstrapping test (Figure 4, also see Supplementary Fig. S7 online). However, the network revealed higher genetic connectivity between Fiq and Jigjiga than Dire Dawa (Figure 4a). This finding suggests Jigjiga is the potential source population for the studied Fiq *An. stephensi*.

**Figure 4.**
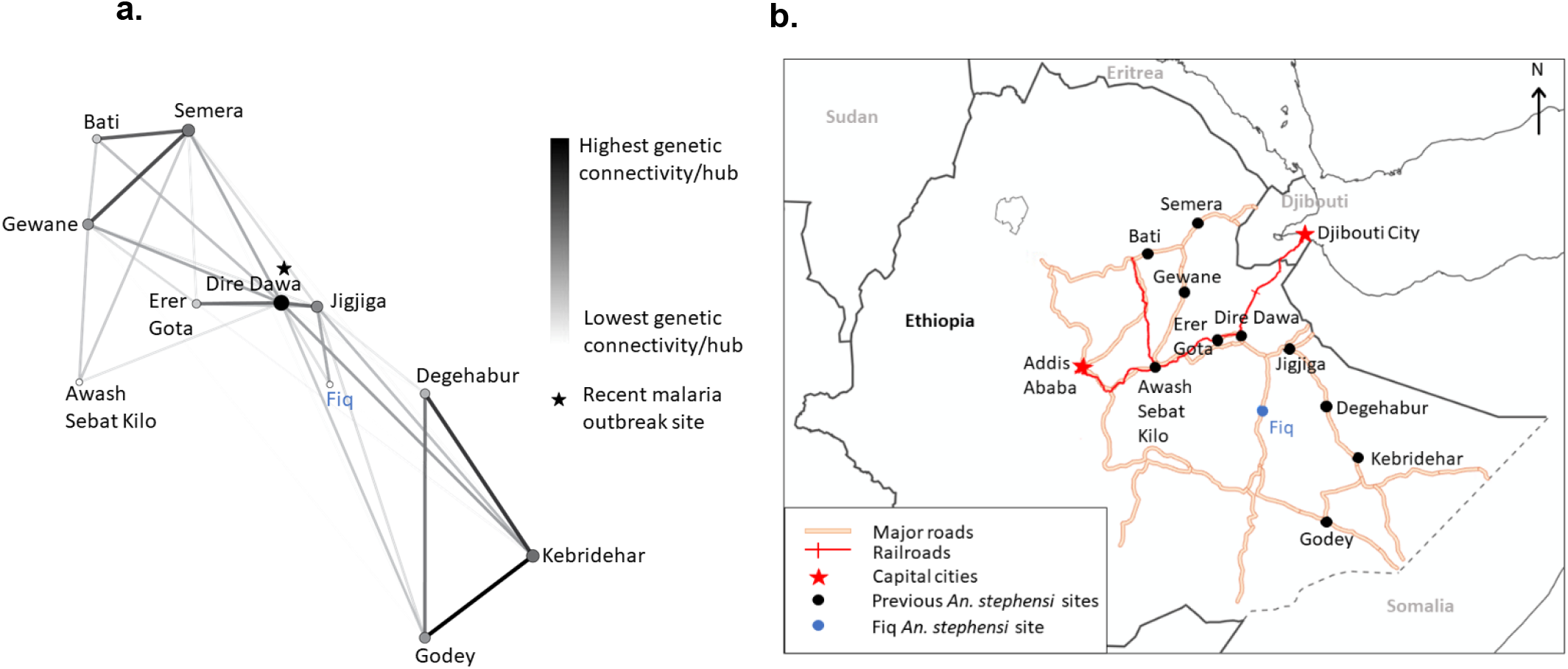
**a**. Genetic network of *An. stephensi* populations in eastern Ethiopia. Network nodes represent populations/hubs and links represent weighted genetic distances/genetic connectivities. The figure is produced by EDENetworks based on a genotype autosomal ddRAD-seq loci matrix by applying a single realization of bootstrapping with 0.85 percentage of nodes at each location and thresholded at 0.12. The colors and sizes of the links represent the strength of genetic connectivity from lowest (white) to highest (black). The colors and sizes of the nodes represent the cumulative weighted links from lowest (white) to highest (black). **b**. Map showing Fiq road connections to other sites. The map was generated using ArcGIS Pro v 3.1 (Environmental Systems Research Institute (ESRI), Redlands, CA, USA).

## Discussion

We report the presence of *An. stephensi* in abundance during a rainy season (May-June 2022) in Fiq, Somali region, Ethiopia. From the more than three thousand and five hundred *Anopheles* larvae collected, all were reared and morphologically identified as *An. stephensi*. Molecular identification of a subset for further molecular analyses also confirms the studied samples as *An. stephensi*. All *An. stephensi* larval habitats identified were artificial breeding sites such as plastic sheet water storages, covered and uncovered cisterns, and barrels which are consistent with other *An. stephensi* larval habitats reported in eastern Ethiopia^45^. The fact that no other *Anopheles* larvae were collected suggests that *An. stephensi* could persist during dry seasons^15^ in Fiq, characteristics typically distinct from *An. arabiensis*, the primary malaria vector in Ethiopia^46,47^. However, in Kenya, *An. stephensi* larvae were found in both artificial containers and a riverbed setting^48^, emphasizing the potential diversity of larval habitats of this invasive *An. stephensi* that is worth noting for future entomological surveillance of this invasive malaria vector in Ethiopia and Africa.

The newly detected Fiq *An. stephensi* were found to be resistant to bendiocarb (carbamate), cyfluthrin, deltamethrin, permethrin, and alphacypermethrin (pyrethroids), but susceptible to pirimiphos-methyl (organophosphate) according to the WHO criteria of 98% - 100% mortality indicating susceptibility^24^. However, restored susceptibility to deltamethrin and permethrin was observed following the synergist bioassay with piperonyl butoxide (PBO), which is a detoxifying enzyme inhibitor that bolsters the effects of pyrethroid insecticides, thus indicating a metabolic insecticide resistance mechanism ^49,50^. This finding is consistent with previous reports of insecticide resistance among the Ethiopian *An. stephensi* populations ^51^ and suggests that PBO plus pyrethroids may be more effective against adult *An. stephensi* in Fiq. Additionally, Fiq *An. stephensi* larvae were found to be susceptible to the diagnostic dose of temephos (0.25 mg/l) with a 100% mortality rate (Table 1), which is consistent with recent larvicide studies conducted on three other *An. stephensi* populations in eastern Ethiopia ^52,53^. Thus, temephos may be an effective larvicide to control *An. stephensi* larvae in Fiq.

Also, our molecular analysis of the pyrethroid target site knockdown gene revealed the presence of the *kdr* mutation in a single pyrethroid-resistant Fiq sample (Figure 2). The low *kdr* mutation frequency among the Fiq pyrethroid-resistant studied *An. stephensi* is consistent with previous low *kdr* mutation frequency in the Ethiopian *An. stephensi* from eastern Ethiopia^27^. This finding confirms that *kdr* mutations may not be a primary mechanism of pyrethroid resistance in the invasive *An. stephensi*^*27*^. Thus, the metabolic resistance mechanism again seems more relevant, as revealed by the positive synergist assays in pyrethroid-resistant *An. stephensi* in this study (Fig.1), and previously reported in Balkew et al.^54^. However, the analysis of gene expression at metabolic resistance loci is needed to fully understand the underlying molecular mechanisms of insecticide resistance development in the invasive *An. stephensi*.

Furthermore, comparative population structure analyses of the newly detected *An. stephensi* with other Ethiopian *An. stephensi* revealed high genetic similarities between Fiq *An. stephensi* and east central *An. stephensi* (Fig. 3b and 3c) and the recentness of Fiq *An. stephensi* compared to them. The recency of Fiq *An. stephensi* was further revealed by their low genetic diversity and high Tajima’s D compared to the east-central Ethiopia *An. stephensi* populations. Thus, these findings suggest that Fiq *An. stephensi* population is from a recent founder event, such as a bottleneck event from east central Ethiopia *An. stephensi* populations. Further population pairwise F_st_ and genetic network analyses pinpoint Jigjiga as the potential source of the founder event in Fiq. Specifically, Fiq *An. stephensi* were less genetically differentiated from Jigjiga *An. stephensi* than the other analyzed Ethiopian *An. stephensi* and shared higher genetic connectivity with Jigjiga than other sites (Figure 4a). Although Fiq is located 195 km away from Jigjiga, the genetic connectivity observed could be explained by the fact that they are connected by a major road (Figure 4b). The same major road also connects both Jigjiga and Fiq to Dire Dawa, the site of a recent *An. stephensi*-mediated malaria outbreak ^7^. Thus, our finding highlights the potential role of roads, perhaps through human and/or good transportation, in the expansion of *An. stephensi* into Fiq. The role that transports routes (marines, roads, etc.) play in invasive species incursion into new geographic areas has been well documented ^55-58^. Thus, this potential mechanism of *An. stephensi* dispersal within the region warrants further investigation.

## Conclusion

The present study revealed the high prevalence of the invasive *An. stephensi* in Fiq, its larval habitats, insecticide resistance status for both adult and immature stages, genetic diversity, population structure, and potential source populations. Our results showed that Fiq *An. stephensi* population is susceptible to pirimiphos-methyl, PBO-pyrethroids, and temephos. Thus, these insecticides may be effectively used in control strategies against this invasive malaria vector in Fiq. We also found that the Fiq *An. stephensi* population shares genetic connectivity with two major *An. stephensi* hubs (i.e., Jigjiga and Dire Dawa) in eastern Ethiopia, with a stronger connection to Jigjiga. Thus, heightened vector control in those areas could help prevent further *An. stephensi* incursions into Fiq and beyond. Overall, this study provides a comprehensive approach to investigate a recent *An. stephensi* expansion into a new geographical area to determine the extent of establishment, assess effectiveness of insecticides, and identify potential source populations to prevent further spread.

## Data Availability Statement

DNA sequences: Bioproject PRJNA1042829 and Genbank accessions SAMN38322421 – SAMN38322440.

## Acknowledgments

We are thankful to the field teams who collected and organized the mosquitoes. We thank Dr. Philip Lavretsky (University of Texas at El Paso) for the Illumina barcodes. We also thank Ms. Madison Follis, Ms. Amelia Wickham, and Ms. Nidhi Kotha (Baylor University) for their lab support and Mr. Joshua Been (Baylor Library Data and Digital Scholarship) for his support with the GIS road map.

## Author’s contribution

JNS, SY, and TEC conceptualized the study. SY, MA, TEC, and SZ organized field collections. SY conducted the bioassays. JNS and TEC collected and analyzed the molecular and genomic data. JNS, SY, SZ, and TEC contributed to the writing of the manuscript. All authors reviewed and approved the final manuscript.

## Disclaimer

The authors alone are responsible for the views expressed in this article and they do not necessarily represent the views, decisions, or policies of the institutions with which they are affiliated.

## Supplementary Information

