## Supplemental Information for "Insecticide resistance and population structure of the invasive malaria vector, *Anopheles stephensi*, from Fiq, Ethiopia"

**Supplementary information**

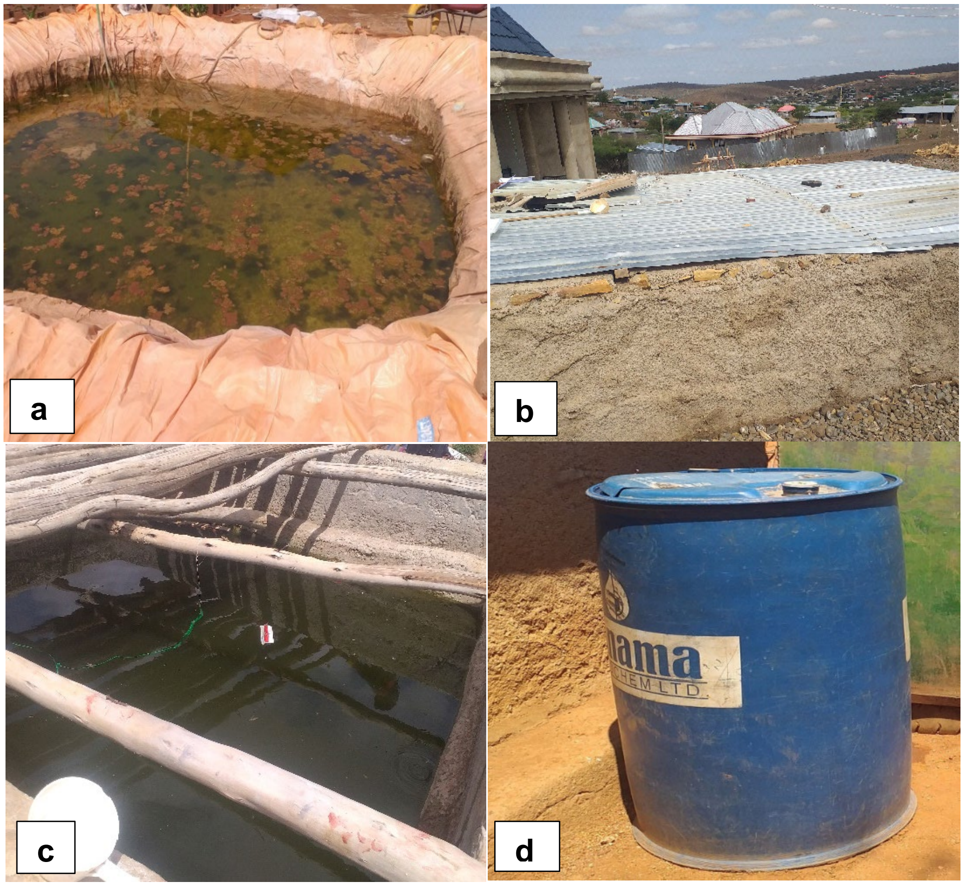


**Figure S1.** Breeding habitat types: **a.** Plastic sheet water storage **b.** Covered cistern **c.** Uncovered cistern **d.** Barrel

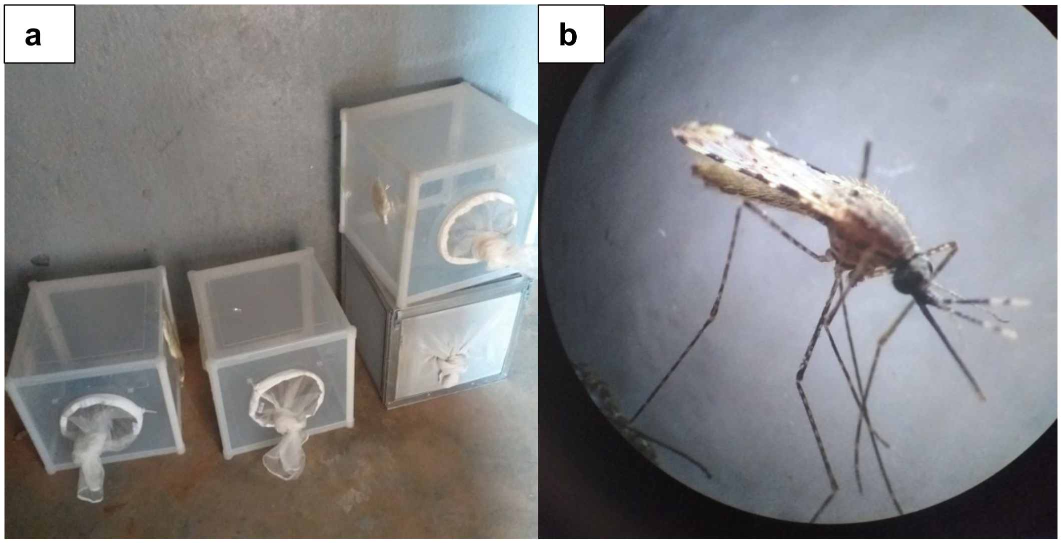


**Figure S2. a.** Rearing of *An. stephensi* at field insectary. **b.** Adult *An. stephensi* reared from wild larvae and pupae.

**Table S1.** Adult mosquito collection

| **Collection methods** | **Fiq** | |
|---|---|---|
| | *An. stephensi* | *Culex* spp. |
| CDC LT | 0 | 123 |
| PSC | 0 | 87 |
| Prokopack (Goat shed) | 2 | 9 |
| Total | 2 | 219 |

**Table S2.** Types of mosquito larval breeding habitats in Fiq town

| Type of container | Number of containers inspected | Number of positive containers | Total larvae | | Number of adult mosquitoes reared from *Anopheles* larvae |
|---|---|---|---|---|---|
| | | | *Anopheles* | *Culex* | *An. stephensi* |
| Cisterns | 17 | 15 | 2066 | 132 | 2066 |
| Plastic containers | 8 | 8 | 1423 | 181 | 1423 |
| Barrels | 6 | 3 | 224 | 77 | 224 |
| Total | 31 | 26 | 3713 | 390 | 3713 |

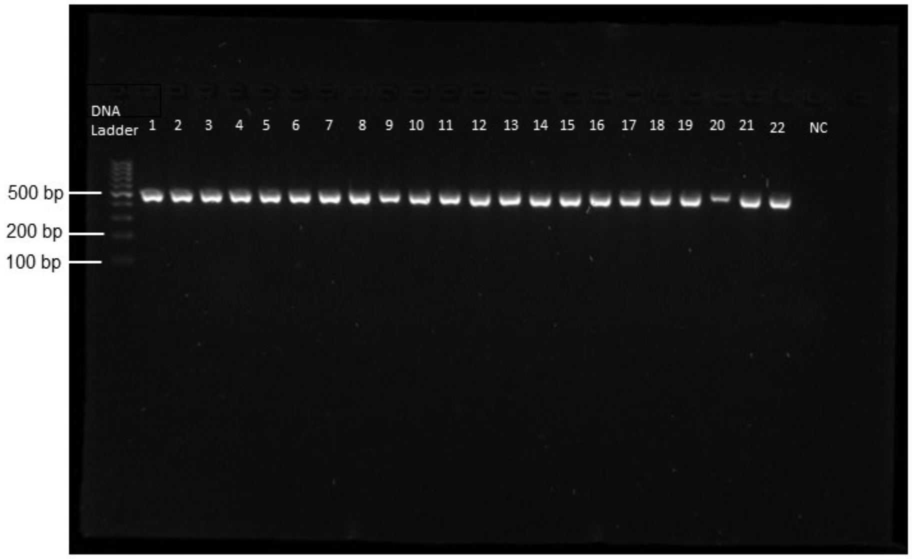


**Figure S3**. Gel photograph showing the result of ITS2 endpoint assay. Lanes 1-22 represent DNA samples from morphologically identified *An. stephensi* from Fiq, Ethiopia. NC stands for negative control.

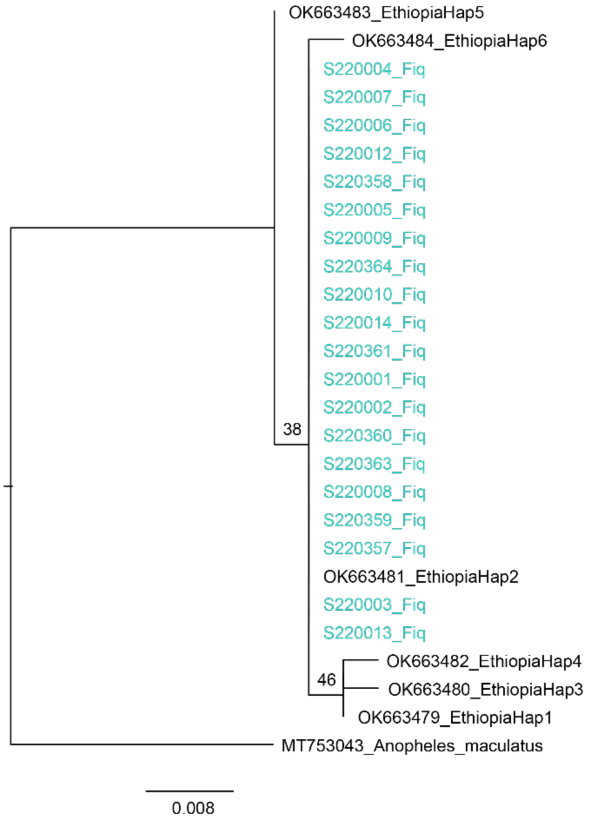


**Figure S4**. Phylogenetic analysis of Ethiopian *An. stephensi* cytochrome oxidase subunit I (COI) sequenced data based on maximum likelihood approach. The tree with the highest likelihood score is shown. Fiq *An. stephensi* sequences are shown in color.

**Table S3.** Mortality rates of *An. stephensi* exposed to different insecticides in Fiq, Ethiopia

| Type of assay | Classification | Insecticides discriminating concentration (%) | No. of *An. stephensi* tested | Mortality rate after 24hrs (%) | | | | Mortality rate (%) mean [95% CI] | Resistance (<90%) |
|---|---|---|---|---|---|---|---|---|---|
| | | | | R1 | R2 | R3 | R4 | | |
| Diagnostic dose | Carbamates | Bendiocarb 0.1% | 100 | 40 | 40 | 32 | 48 | 40 [30, 50] | Yes |
| | Pyrethroid | Cyfluthrin 0.15% | 100 | 96 | 76 | 88 | 88 | 87 [74, 100] | Yes |
| | | Deltamethrin 0.05% | 100 | 56 | 64 | 64 | 80 | 66 [50, 82] | Yes |
| | | Permethrin 0.75% | 100 | 52 | 60 | 64 | 76 | 63 [47, 79] | Yes |
| | | Alphacypermethrin 0.05% | 100 | 76 | 52 | 40 | 16 | 46 [6, 86] | Yes |
| | Organophosphorus | Pirimiphos-methyl 0.25% | 100 | 100 | 100 | 100 | 100 | 100 | No |
| Synergist assays | Pyrethroid | Deltamethrin 0.05% | 75 | 88 | 28 | 48 | - | 55 [-21, 130] | Yes |
| | | Deltamethrin + PBO | 75 | 100 | 100 | 100 | - | 100 | No |
| | | Permethrin 0.75% | 75 | 84 | 80 | 48 | - | 71 [22, 119] | Yes |
| | | Permethrin + PBO | 75 | 100 | 100 | 96 | - | 99 [99, 104] | No |

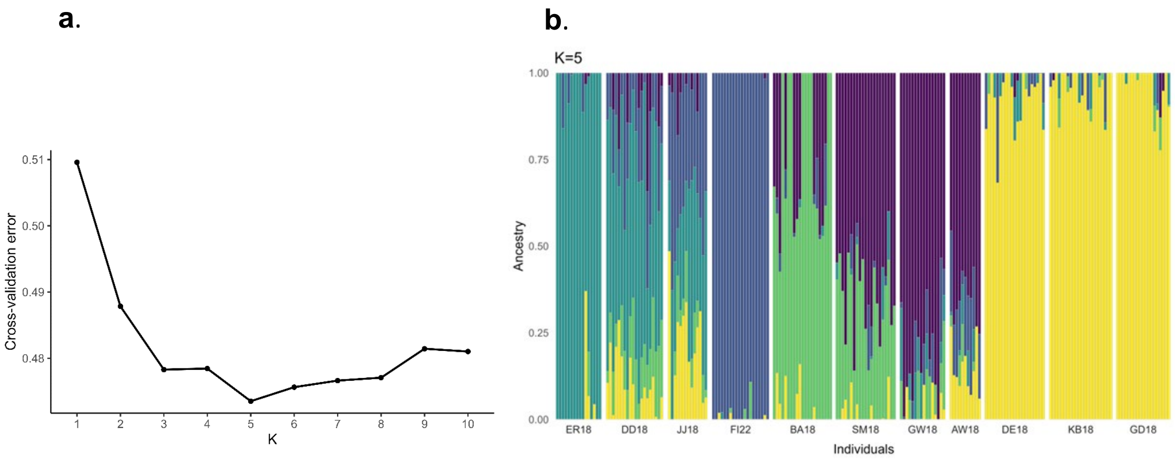


**Figure S5. a.** Admixture cross-validation errors of K values. **b.** Ancestry plot based on ADMIXTURE analysis. Each barplot represents an individual. The height of each colored bar represents the probability of assignment to the corresponding ancestry when assuming the presence of five ancestral populations (K = 5). The sample sites are Erer Gota (ER18), Dire Dawa (DD18), Jigjiga (JJ18), Fiq (FI22), Bati (BA18), Semera (SM18), Gewane (GW18), Awash Sebat Kilo (AW18), Degehabur (DE18), Kebriderar (KB18), and Godey, (GD18).

| | | Northeastern Ethiopia | | | | East-central Ethiopia | | | Southeastern Ethiopia | | | Fiq |
|---|---|---|---|---|---|---|---|---|---|---|---|---|
| | | Semera | Bati | Gewane | Awash Sebat Kilo | Erer Gota | Dire Dawa | Jigjiga | Degehabur | Kebridehar | Godey | Fiq |
| Northeastern Ethiopia | Semera | . | 0.05 | 0.05 | 0.07 | 0.10 | 0.06 | 0.08 | 0.12 | 0.11 | 0.10 | 0.12 |
| | Bati | 0.05 | . | 0.08 | 0.10 | 0.13 | 0.08 | 0.10 | 0.14 | 0.14 | 0.14 | 0.14 |
| | Gewane | 0.05 | 0.08 | . | 0.06 | 0.09 | 0.06 | 0.08 | 0.11 | 0.11 | 0.10 | 0.12 |
| | Awash Sebat Kilo | 0.07 | 0.10 | 0.06 | . | 0.10 | 0.07 | 0.08 | 0.10 | 0.10 | 0.10 | 0.10 |
| East-central Ethiopia | Erer Gota | 0.10 | 0.13 | 0.09 | 0.10 | . | 0.06 | 0.08 | 0.12 | 0.12 | 0.12 | 0.12 |
| | Dire Dawa | 0.06 | 0.08 | 0.06 | 0.07 | 0.06 | . | 0.04 | 0.09 | 0.08 | 0.08 | 0.08 |
| | Jigjiga | 0.08 | 0.10 | 0.08 | 0.08 | 0.08 | 0.04 | . | 0.08 | 0.08 | 0.08 | 0.07 |
| Southeastern Ethiopia | Degehabur | 0.12 | 0.14 | 0.11 | 0.10 | 0.12 | 0.09 | 0.08 | . | 0.04 | 0.05 | 0.11 |
| | Kebridehar | 0.11 | 0.14 | 0.11 | 0.10 | 0.12 | 0.08 | 0.08 | 0.04 | . | 0.04 | 0.11 |
| | Godey | 0.10 | 0.14 | 0.10 | 0.10 | 0.12 | 0.08 | 0.08 | 0.05 | 0.04 | . | 0.13 |
| Fiq | Fiq | 0.12 | 0.14 | 0.12 | 0.10 | 0.12 | 0.08 | 0.07 | 0.11 | 0.11 | 0.13 | . |

**Figure S6.** Population pairwise $F_{st}$ based on autosomal loci. Color gradient based on $F_{st}$ values (yellow = highest, dark orange = lowest).

**Table S4.** Basic summary statistics based on autosomal loci.

| Collection site | Sample size N | Nucleotide diversity π | Counts of segregating SNPs S | Tajima's D |
|---|---|---|---|---|
| **Fiq Ethiopia** | | | | |
| Fiq | 20 | 0.1986 | 1071 | 1.29 |
| **Northeastern Ethiopia** | | | | |
| Semera | 21 | 0.2255 | 1374 | 0.76 |
| Bati | 21 | 0.2084 | 1218 | 0.95 |
| Gewane | 16 | 0.2225 | 1291 | 0.71 |
| Awash Sebat Kilo | 11 | 0.2300 | 1211 | 0.74 |
| **East-central Ethiopia** | | | | |
| Erer Gota | 16 | 0.2195 | 1197 | 0.97 |
| Dire Dawa | 20 | 0.2261 | 1366 | 0.70 |
| Jigjiga | 14 | 0.2107 | 1198 | 0.66 |
| **Southeastern Ethiopia** | | | | |
| Degehabur | 21 | 0.2018 | 1084 | 1.36 |
| Kebridehar | 22 | 0.2087 | 1101 | 1.48 |
| Godey | 18 | 0.1979 | 1078 | 1.19 |

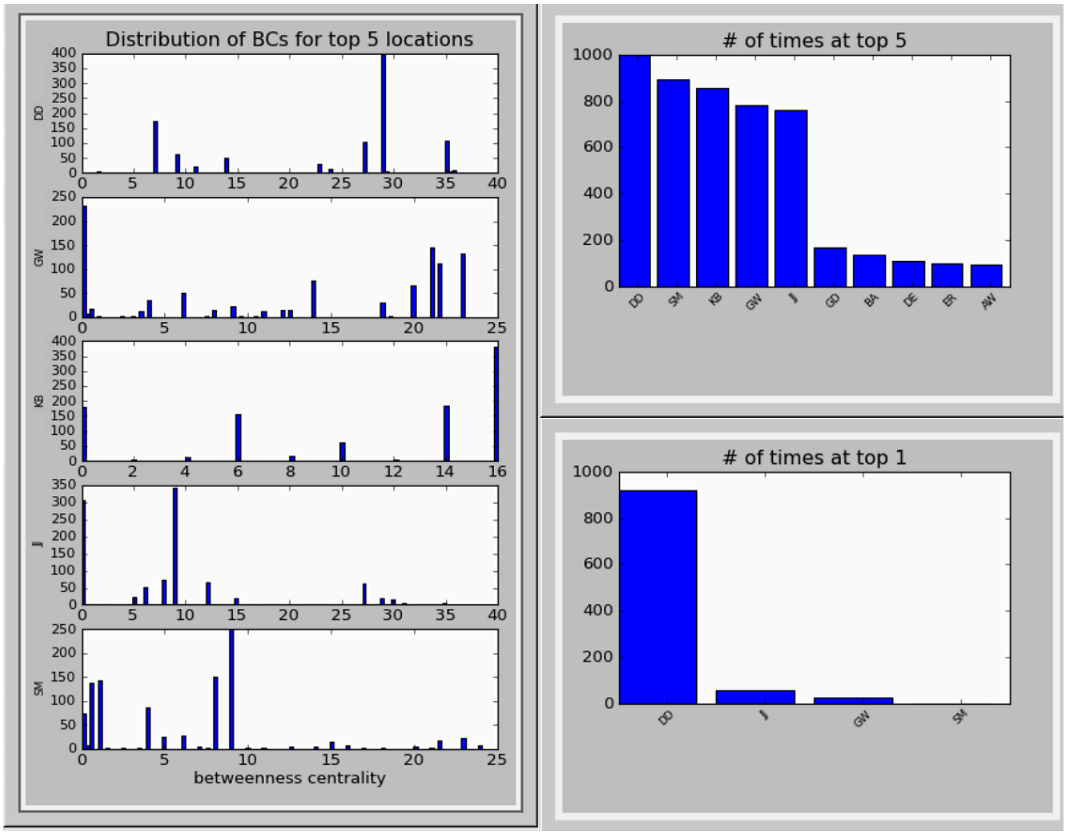


**Figure S7.** Bootstrapping results for betweenness centrality with number of repetitions (1000) and threshold set at 0.85 for Fiq, Ethiopia and eastern Ethiopia *An. stephensi* genetic networks.

**Figure S8.** Alignment of *kdr* locus indicating *kdr* mutation L1014F

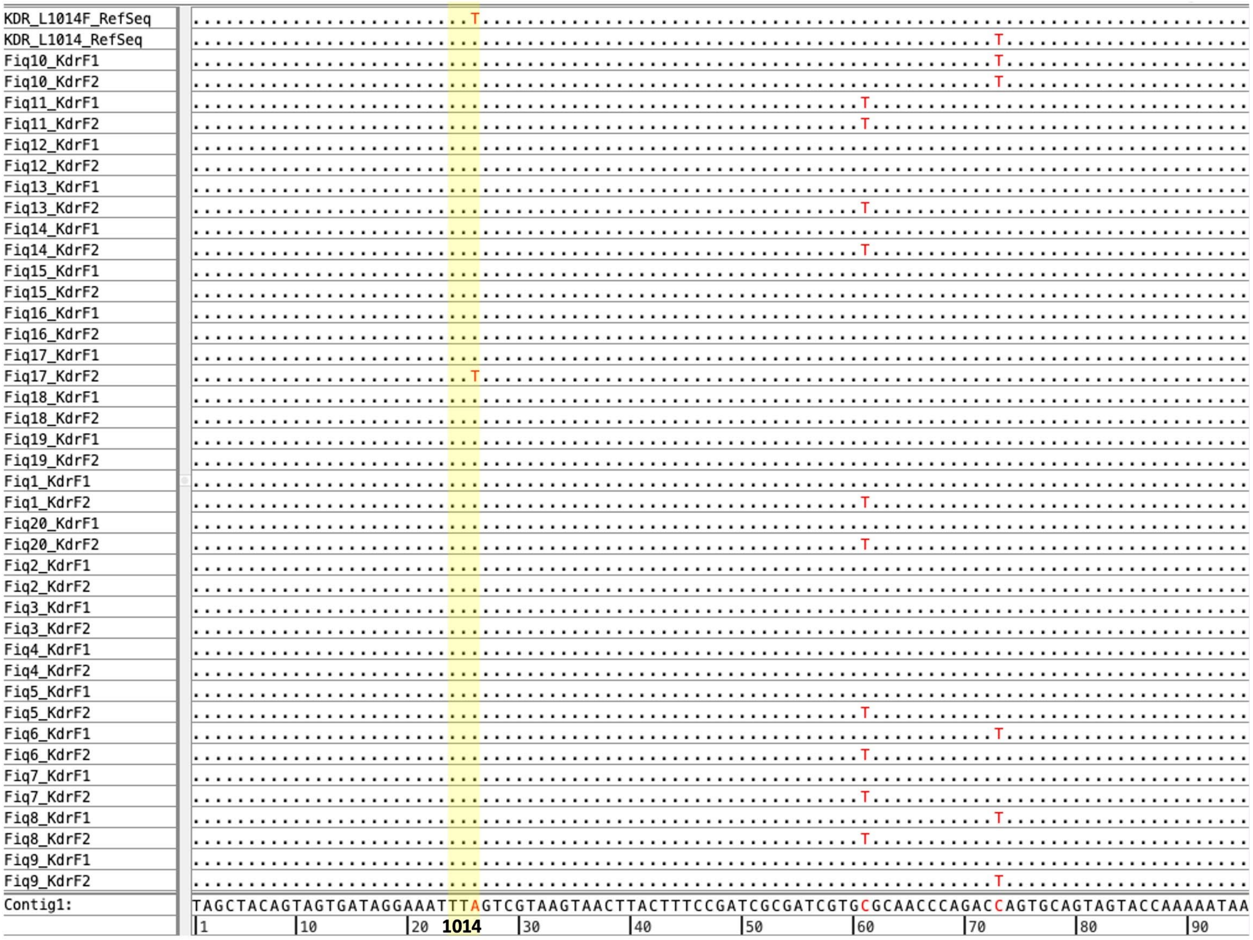

**Table S5.** Fiq *An. stephensi kdr* mutation status.

| Fiq samples | | Insecticide | Status | KDR mutation/variant |
|---|---|---|---|---|
| 1. | S220007 | Deltamethrin | Alive | No |
| 2. | S220008 | Deltamethrin | Alive | No |
| 3. | S220009 | Deltamethrin | Dead | No |
| 4. | S220010 | Deltamethrin | Dead | No |
| 5. | S220012 | Permethrin | Alive | No |
| 6. | S220013 | Permethrin | Dead | No |
| 7. | S220014 | Permethrin | Dead | No |
| 8. | S220357 | Alphacypermethrin | Dead | No |
| 9. | S220358 | Alphacypermethrin | Dead | No |
| 10. | S220359 | Alphacypermethrin | Alive | Yes/heterozygous L1014F |
| 11. | S220360 | Alphacypermethrin | Alive | No |
| 12. | S220361 | Cyfluthrin | Dead | No |
| 13. | S220363 | Bendiocarb | Dead | No |
| 14. | S220364 | Bendiocarb | Alive | No |
| 15. | S220005 | Pirimiphos-methyl | Dead | No |
| 16. | S220006 | Pirimiphos-methyl | Dead | No |
| 17. | S220001 | PBO+Deltamethrin | Dead | No |
| 18. | S220002 | PBO+Deltamethrin | Dead | No |
| 19. | S220003 | PBO+Permethrin | Dead | No |
| 20. | S220004 | PBO+Permethrin | Dead | No |

**Supplementary Document S1**

ddRAD-seq Library Preparation Methods

Genome-wide single nucleotide polymorphism (SNP) data was collected using the double-digest restriction-site associated DNA sequencing (ddRADseq) protocol outlined in Lavretsky et al. (2015), but with fragment size selection following Hernandez et al. (2021). A total of 20 samples with ~20 ng of genomic DNA each were enzymatically fragmented using 1 µL each of SbfI and EcoRI restriction enzymes. Illumina TruSeq compatible adapters and 6 base-pair barcodes were ligated to allow for future de-multiplexing. The adapter-ligated DNA fragments were then size selected using a double-sided size selection based on a total of 0.8x solution of sparQ PureMag beads (Quantabio, MA, USA). First, a right-sided selection for large fragments was completed by adding 0.55x concentration of the total starting ligated DNA solution. The solution was incubated at room temperature for 10 minutes, then transferred to a magnetic plate to rest for 5 minutes or until the mixture became clear. The supernatant containing both target-sized and small-sided DNA fragments was then transferred to new tubes and the beads were discarded. Next, a left-sided size selection against small DNA fragments (<100bp) was completed by adding 0.25x concentration of the total starting ligated DNA solution. The solution was again incubated at room temperature for 10 minutes, then transferred to a magnetic plate to rest for 5 minutes at room temperature or until the mixture became clear. The supernatant containing small-sided DNA fragments (<100bp) was then discarded and the beads were washed twice with 70% ethanol. The beads were then air-dried at room temperature for 2-5 minutes. DNA was re-suspended with 22 µL molecular grade water and eluted for 1 hour, then transferred to a magnetic plate to rest for 1 minute at room temperature or until the mixture became clear. The eluant was then transferred to new tubes. Target size selected DNA fragments were then PCR amplified with Phusion High-Fidelity DNA polymerase, and 10x concentration of forward and reverse RAD primers (Dacosta & Sorenson, 2014) under the following PCR conditions: an initial 30 sec cycle at 98 $^{o}$C, followed by 22 cycles of 10 sec at 98 $^{o}$C, 30 sec at 60 $^{o}$C, and 40 sec at 72 $^{o}$C, with a final extension at 72 $^{o}$C for 5 min. Amplicons were then cleaned using a 1.8x solution of sparQ PureMag beads (Quantabio, MA, USA) and two 70% ethanol washed before final elution in 40 µL molecular grade water. Library concentrations were quantified using Qubit dsDNA BR Assay Kit (ThermoFisher Scientific, MA, USA) following manufacturer protocols. Samples were then pooled in equimolar amounts and the multiplexed library was sequenced on Illumina Hiseq X using single-end 150 bp chemistry with Novogene (Novogene Inc., CA, USA).
